# Mitochondrial fission is a critical modulator of mutant APP-induced neural toxicity

**DOI:** 10.1101/2020.05.10.087346

**Authors:** Lauren Y. Shields, Huihui Li, Kevin Nguyen, Hwajin Kim, Zak Doric, T. Michael Gill, Dominik Haddad, Keith Vossel, Meredith Calvert, Ken Nakamura

## Abstract

Alterations in mitochondrial fission may contribute to the pathophysiology of several neurodegenerative diseases, including Alzheimer’s disease (AD). However, we understand very little about the normal functions of fission, or how fission disruption may interact with AD-associated proteins to modulate pathogenesis. Here we show that loss of the central mitochondrial fission protein dynamin-related 1 (Drp1) in CA1 and other forebrain neurons markedly worsens the learning and memory of mice expressing mutant human amyloid-precursor protein (hAPP) in neurons. In cultured neurons, Drp1KO and hAPP converge to produce mitochondrial Ca^2+^ (mitoCa^2+^) overload, despite decreasing mitochondria-associated ER membranes (MAMs) and cytosolic Ca^2+^. This mitoCa^2+^ overload occurs independently of ATP levels. These findings reveal a potential mechanism by which mitochondrial fission protects against hAPP-driven pathology.

## INTRODUCTION

Although the pathophysiology of Alzheimer’s disease (AD) remains poorly understood, several lines of evidence suggest that mitochondrial dysfunction and disrupted balance between mitochondrial fission and fusion may contribute to neurodegeneration (1). Indeed, levels of the central mitochondrial fission protein dynamin-related protein 1 (Drp1) are frequently increased in post-mortem tissue from AD patients, while fusion proteins are decreased (2,3), suggesting a shift towards fission. In addition, mutations or overexpression of a number of disease-related proteins (4–8), including amyloid precursor protein (APP) (9) and amyloid beta (Aβ) (10), augment mitochondrial fragmentation, and Aβ has been proposed to cause toxicity by increasing function of Drp 1 (10). Although it remains unclear whether increased fragmentation actually causes neurodegeneration, inhibiting Drp1 can protect against toxicity in models of Parkinson’s disease and Huntington’s disease (7,8,11,12).

Importantly, disrupting the fission-fusion balance in either direction can be toxic, since insufficient mitochondrial fission produces excessive mitochondrial tubulation and also causes disease (1). Mutations in Drp1 are now recognized to cause a range of neurologic disorders from encephalopathy and neonatal lethality to refractory epilepsy (13–16). In addition, tau can cause toxicity by blocking Drp1 localization to mitochondria (17). Therefore, the role of Drp 1 in AD pathogenesis still remains unclear and may be contextdependent.

The functions of mitochondrial fission in healthy neurons also remain poorly understood, but appear to be multifactorial. Neurons require Drp1 to target mitochondria down distal axons (18,19). In addition, neurons lacking Drp1 are unable to maintain normal levels of mitochondrial-derived ATP at the nerve terminal (20), at least in part due to intrinsic functional deficits in mitochondria lacking Drp1. Drp1 is also critical for respiration in other cells with high energy requirements such as cardiac myocytes (21), whereas the effect of Drp1 loss in other cell types, such as mouse embryonic fibroblasts (MEFs), is less consistent (20–27). Fission is also widely hypothesized to support mitochondrial function by facilitating the turnover of dysfunctional mitochondria (28). Without fission, mitochondria may be too large to be engulfed by autophagosomes, causing dysfunctional mitochondria to accumulate (20,22). However, mitochondrial fission may also support mitochondrial function more directly. For instance, mitochondrial fission by Drp1 occurs at the points of contact between mitochondria and the ER (mitochondria-associated membranes, MAMs (29)), raising the possibility that Drp1 influences MAM function, and thus Ca^2+^ and lipid metabolism. Interestingly, cytosolic calcium (cytCa^2+^) promotes Drp1 recruitment to mitochondria through calcineurin and Ca^2+^/calmodulin-dependent protein kinase 1 alpha (30,31), although it remains unclear how Drp1 ultimately impacts calcium homeostasis, and whether any such effects are independent of its role in supporting ATP levels in neurons (20).

In order to better understand the functions of mitochondrial fission and how altered fission may influence the pathophysiology of AD, we examined the impact of loss of mitochondrial fission on the toxicity of mutant hAPP in mouse neurons.

## RESULTS

### Loss of mitochondrial fission increases the toxicity of mutant APP in vivo

Increased mitochondrial fission has been proposed to underlie Aβ toxicity (32), and Drp1 inhibitors are protective in other mouse models of neurotoxicity (11,33). To test if loss of the central mitochondrial fission protein Drp 1 protects against the toxicity of mutant APP, we generated mice with targeted deletion of Drp1 on a mutant human APP background. First, Drp1^lox/lox^ mice were crossed with an Alzheimer’s mouse model, hAPP-J20 (34), referred to henceforth as hAPP, which expresses mutant hAPP (Swedish and Indiana mutations) in neurons under the PDGF-beta promoter. All hAPP mice were heterozygous carriers of the hAPP transgene. hAPP;Drp1^wt/lox^ were bred to homozygosity (hAPP;Drp1^lox/lox^). To generate hAPP mice that lack Drp1 in the CA1 and other forebrain neurons (Drp1cKO), we bred CamKCre;Drp1^wt/lox^ and hAPP;Drp1^lox/lox^ mice to generate hAPP;Drp1^lox/lox^;CamKII-Cre mice. The resulting progeny (total n=246 mice) were born in roughly Mendelian proportions including controls that lacked Cre (Drp1^wt/lox^ and Drp1^lox/lox^, 26.4%), Drp1 heterozygotes (Drp1^lox/wt^;CamKII-Cre,13.0%), Drp1cKO (Drp1^lox/lox^;CamKII-Cre, 16.7%), hAPP mice that lacked Cre (hAPP;Drp1^wt/lox^ and hAPP;Drp1^lox/lox^, 20.3%), hAPP Drp1 heterozygotes 13.8%, and hAPP Drp1cKO (9.75%).

As expected, hAPP mice had decreased survival (Fig. 1*A*) (35). Loss of Drp1 expression did not affect the survival defect in hAPP mice, and all genotypes had similar body weights through 7 months of age (Fig. 1*B*). In open field tests, Drp1cKO, hAPP and hAPPDrp1cKO all displayed increased movement in the open field (Fig. 1*C*), consistent with hyperactivity previously seen in hAPP mice (36) and indicating intact motor activity, as expected for the hAPP genotype. All genotypes exhibited similar swim speeds throughout procedural and spatial learning (Fig. 1*D*), and intact procedural learning during cued platform testing using the Morris water maze (MWM), indicating intact sensorimotor function. Although procedural learning was attenuated in the hAPP and Drp1cKO groups, all groups except for hAPP Drp1cKO mice were equivalent in their procedural performance at the end of training (Fig. 1*E*). However, rather than protecting against hAPP toxicity, Drp1KO markedly exacerbated the spatial learning and memory impairments of hAPP mice in the MWM. hAPP Drp1cKO mice were unable to learn the platform location despite training over 14 sessions during 7 consecutive days (Fig. 1*F*), indicating a strong functional synergism of Drp1 loss and hAPP in vivo. Drp1cKO and hAPP mice showed only subtle spatial learning deficits, based on rank order analysis of latency. No learning differences were noted with Drp1 heterozygotes (Fig. S1A, S1B). Spatial learning and memory deficits of Drp1cKO and hAPP mice were confirmed during probe trial performance carried out 24 and 72 hours after the last training trial. hAPP mice took significantly longer to cross over the former location of the hidden platform (target), and both Drp1cKO and hAPP mice did not cross its former location as frequently as control mice. hAPP Drp1cKO mice performed worse than mice harboring either mutation alone (Fig. 1*G* and *H*). Therefore, complete Drp1 loss markedly worsens (rather than prevents) the adverse effects of hAPP on memory. In contrast, hAPP Drp1cHET mice had decreased latency to target at 72 hours compared to hAPP mice, and also crossed the former platform location more frequently than hAPP mice at 24 hours. Therefore, partial Drp1 loss does not worsen spatial learning and memory, and may actually protect against it (Fig. S1C, S1D).

**Figure 1.**
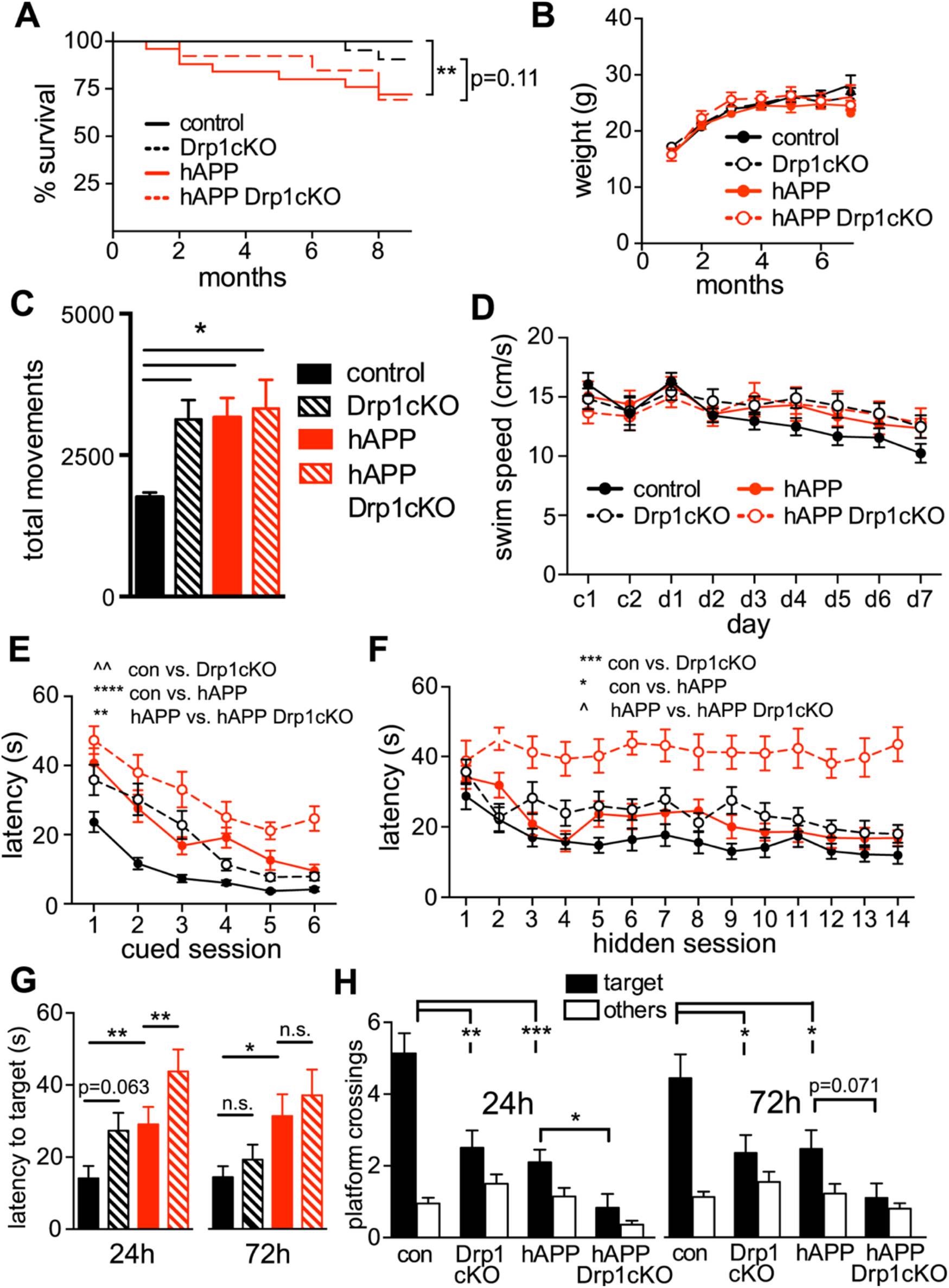
Drp1 loss and hAPP expression synergistically impair learning and memory. *A*, hAPP (hAPP-J20;Drp1^wt/lox^ and hAPP-J20;Drp1^lox/lox^) mice had significant premature mortality (**p<0.01) compared to controls (Drp1^wt/lox^ and Drp1^lox/lox^); hAPP Drp1cKO (hAPP-J20;Drp1^lox/lox^;CamKII-Cre) mice had a trend (p=0.11) towards premature mortality compared to Drp1cKO (Drp1^lox/lox^;CamKII-Cre), by Log-rank Mantel-Cox test, n=13-30 mice/group monitored from birth through 9 months of age. *B*, No weight differences were observed between genotypes up to 7 months. Data are means ± S.E.M.; n=5-42 mice/group. *C*, 6–7-month-old Drp1cKO, hAPP, and hAPP Drp1cKO mice showed an increased number of total movements in an open field over the course of 15 min, as compared to controls (Drp1^wt/lox^ and Drp1^lox/lox^). Data are means ± S.E.M.; *p<0.05 by one-way ANOVA and Holm-Sidak *post hoc* test, n=9-12 mice/group. *D-F,* Both procedural and spatial learning and memory were evaluated using the Morris water maze (MWM). D, No difference in swim speeds were found throughout the 2 days of procedural learning or 7 days of spatial learning by two-way ANOVA with repeated measures, indicating that all groups had intact motor function prior to the start of spatial training. *E*, Procedural cued training conducted on the first 2 days (c1, c2) demonstrated significant but differential learning effects between the groups. Data are means ± S.E.M.; **p<0.01, ****p<0.0001, ^^p<1e-10 by average rank latency with mixed effect modeling, n=12–22 mice/group. *F*, Spatial learning and memory during hidden-platform training in 6–7-month-old mice. Drp1cKO and hAPP showed significant learning impairments compared to Drp1WT (control). hAPP-J20 Drp1cKO (hAPP Drp1cKO) mice showed significant learning impairments compared with hAPP-J20 (hAPP) mice. Data are means ± S.E.M.; *p<0.05, ***p<0.001, ^p<1e-9 by average rank latency with mixed effect modeling, n=12-22 mice/group. *G, H,* Spatial memory was evaluated using MWM probe trials at 24 and 72 h with the hidden-platform removed, and measured by latency to cross the former hidden platform location (target) (G) and number of platform location (target) and non-target (other) crossings (H). Drp1cKO, hAPP, and hAPP Drp1cKO mice showed significant memory deficits. n=12-22 mice/group. Data are means ± S.E.M.; n.s. = not significant, *p<0.05, **p<0.01, ***p<0.001 by Cox proportional hazards regression models (latency to cross) and Quasi-Poisson generalized linear models (platform crossings).

To gain insight into if Drp1 loss increased *in vivo* neurotoxicity by increasing Aβ levels, we examined Aβ deposition. However, concurrent loss of Drp1 did not significantly change the extent of age-dependent Aβ plaque deposition in hAPP mice, although the sensitivity of this experiment was limited by high variability (Fig. S2, *A* and *B*). Likewise, hAPP expression alone did not affect hippocampal and CA1 volume, and also did not contribute to the age-dependent loss of hippocampal or CA1 volume evident between 6 and 12-month old Drp1cKO mice (Fig. 2, *A*-C, Fig. S2, *C* and *D*)(20), or affect CA1 cell density (Fig. *2D* and S2*D*). Therefore, concurrent loss of Drp1 and hAPP did not increase the mild, age-dependent neuronal loss seen in Drp1cKO mice(20).

**Figure 2.**
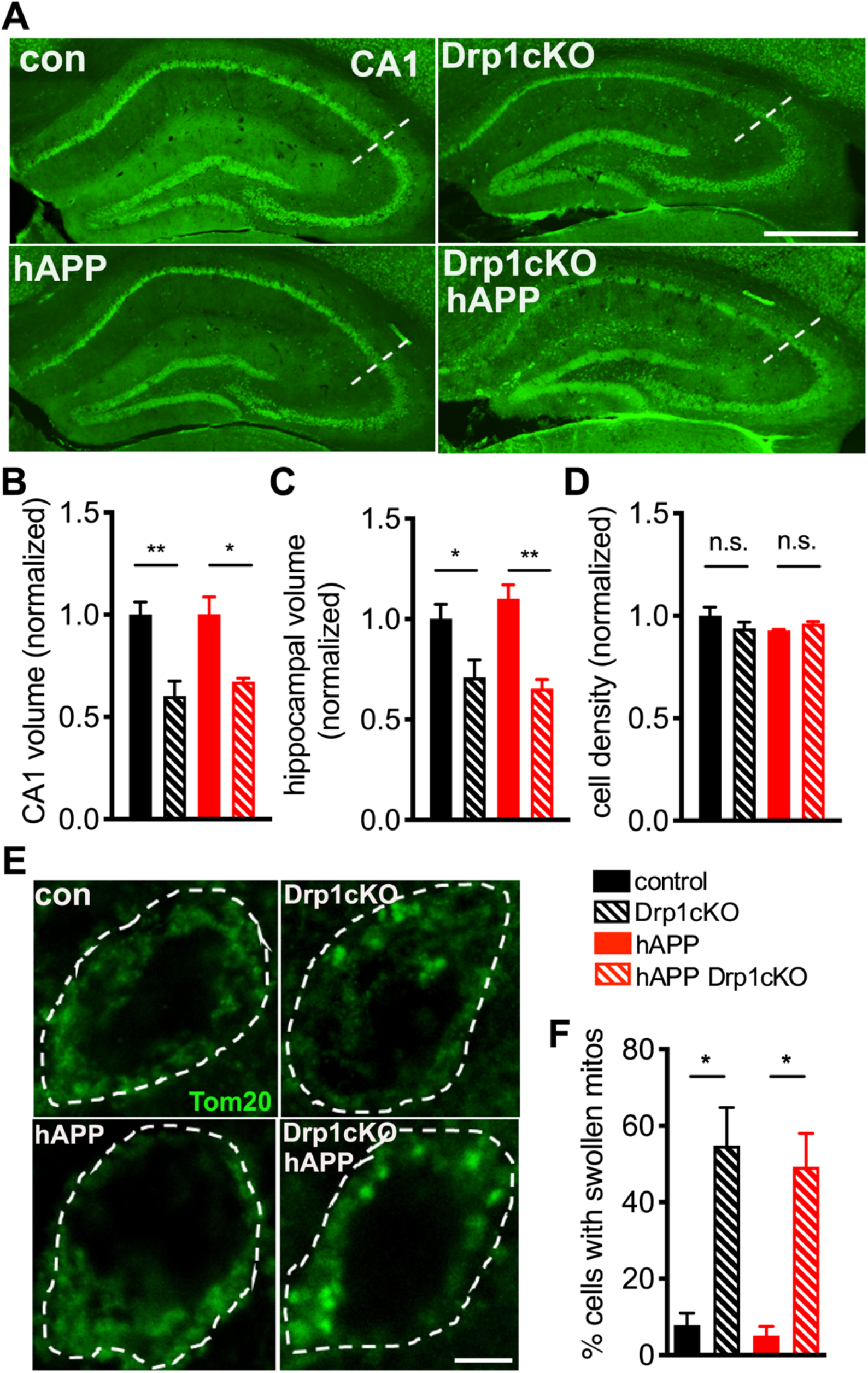
hAPP does not exacerbate Drp1cKO-induced cell loss and morphologic changes. *A*, Neuronal cell bodies labeled by NeuN staining in brain sections from 12-month-old mice. Hippocampi indicated by dotted outlines. Scale bar is 1 mm. *B,C* Drp1 loss decreased both CA1 (B) and overall hippocampal (C) volume in 12-month-old mice. n=4-5 mice/group (9-17 slices/mouse). Data are means ± S.E.M., *p<0.05, **p<0.01 by two-way ANOVA and Holm-Sidak *post hoc* test. *D*, Drp1cKO and hAPP Drp1cKO mice did not show any decrease in CA1 cell density at 12 months of age. n=4-5 mice/group (4 slices/mouse). n.s. (not significant) by two-way ANOVA and Holm-Sidak *post hoc* test. *E*, Mitochondria in CA1 neurons in hippocampal slices from 6-7-month-old Drp1WT (control), Drp1cKO, hAPP-J20 (hAPP), and hAPP-J20 Drp1cKO (hAPP Drp1cKO) mice, identified by Tom20 immunofluorescence (green). Cell bodies (outer stippled outlines) were defined by Map2 staining. Scale bar is 4 μm. *F*, Drp1KO increased the proportion of cells with swollen mitochondria, while hAPP had no effect. n=4 mice/group (3 slices/mouse). *p<0.05 by Welch’s ANOVA and Games-Howell *post hoc* test (used instead of two-way ANOVA due to significant Levene’s test for equality of variance).

### Drp1 KO and hAPP converge to produce mitochondrial Ca^2+^ overload

We next investigated how loss of mitochondrial fission predisposes neurons to the toxicity of hAPP. Drp1KO neurons have swollen mitochondria, and tau may cause toxicity by producing excessive mitochondrial tubulation (17). However, hAPP alone had no impact on mitochondrial morphology, nor did it affect the percent of cells with swollen mitochondria produced by Drp1cKO in vivo (Fig. 2, *E* and *F*). In addition, neither hAPP nor Drp1cKO affected mitochondrial content in hippocampal neurons, as assessed by Tom20 immunofluorescence (Fig. S2*F*), indicating that a change in mitochondrial content is also unlikely to underlie the synergistic toxicity.

Mitochondria have critical functions in Ca^2+^ buffering, which influence both cytosolic and mitochondrial Ca^2+^ (mitoCa^2+^) levels (37,38). Indeed, sufficient mitoCa^2+^ is required for respiratory chain enzyme function, but excessive Ca^2+^ can be toxic. To determine if Drp1KO and hAPP influence mitoCa^2+^ levels, we established a hippocampal neuron model system in which either Drp1 was deleted (Drp1KO), hAPP was overexpressed, or both. Specifically, we co-transfected Drp1^lox/lox^ primary hippocampal neurons with hAPP (39,40) and either Cre recombinase (to remove Drp1) or a vector control, as well as Cepia3mt to measure mitoCa^2+^ (41) and mApple as a control to normalize for probe expression level. We confirmed hAPP expression (Fig. S3*A*), and that Cre expression led to the expected altered mitochondrial morphology and distribution *in vitro,* indicative of Drp1 deletion (data not shown) (20).

We first examined basal levels of mitoCa^2+^, estimated based on the ratio of basal Cepia3mt fluorescence/mApple fluorescence, and found they were similar in all groups (Fig. S3*B*). Next, we examined the neuron’s capacity to buffer Ca^2+^ during neural activity, using electrical field stimulation (30hz for 3s), which promotes preferential release of vesicles in the readily releasable pool (42) (Fig. 3 *A-D*, Fig. S3*C*). Following each electrical pulse train, mitoCa^2+^ transiently increased and then returned to baseline in most cells from all groups (e.g. Fig. 3, *A* and *B*). However, in a subset of cells of all genotypes, mitoCa^2+^ levels failed to recover (i.e. return to <30% peak amplitude) within ≈17s after electrical stimulation, defined as mitoCa^2+^ overload (e.g. Fig. 3*C*). Overall, the combination hAPP-Drp1KO increased both the extent of mitoCa^2+^ influx (Fig. *3D,E*) and the frequency of mitoCa^2+^ overload (Fig. 3*F*), while neither hAPP or Drp1KO alone affected these parameters. The increased mitoCa^2+^ influx was driven by increased mitoCa^2+^ influx into those cells that underwent mitoCa^2+^ overload, since mitoCa^2+^ influx was unchanged among those cells that recovered (Fig. S3*C,D*). Moreover, mitoCa^2+^ overload appears to be triggered by excessive mitoCa^2+^ influx not decreased efflux. Indeed, among cells that recovered, both Drp1KO and hAPP-Drp1KO actually had a shorter half-life of mitoCa^2+^ decay (Fig. S3*E*), raising the possibility that loss of Drp1 upregulates mitoCa^2+^ efflux, for instance to cope with increased import. However, in hAPP-Drp1KO cells, our data suggest that increased mitoCa^2+^ influx eventually exceeds the capacity for mitoCa^2+^ export, resulting in mitoCa^2+^ overload.

**Figure 3.**
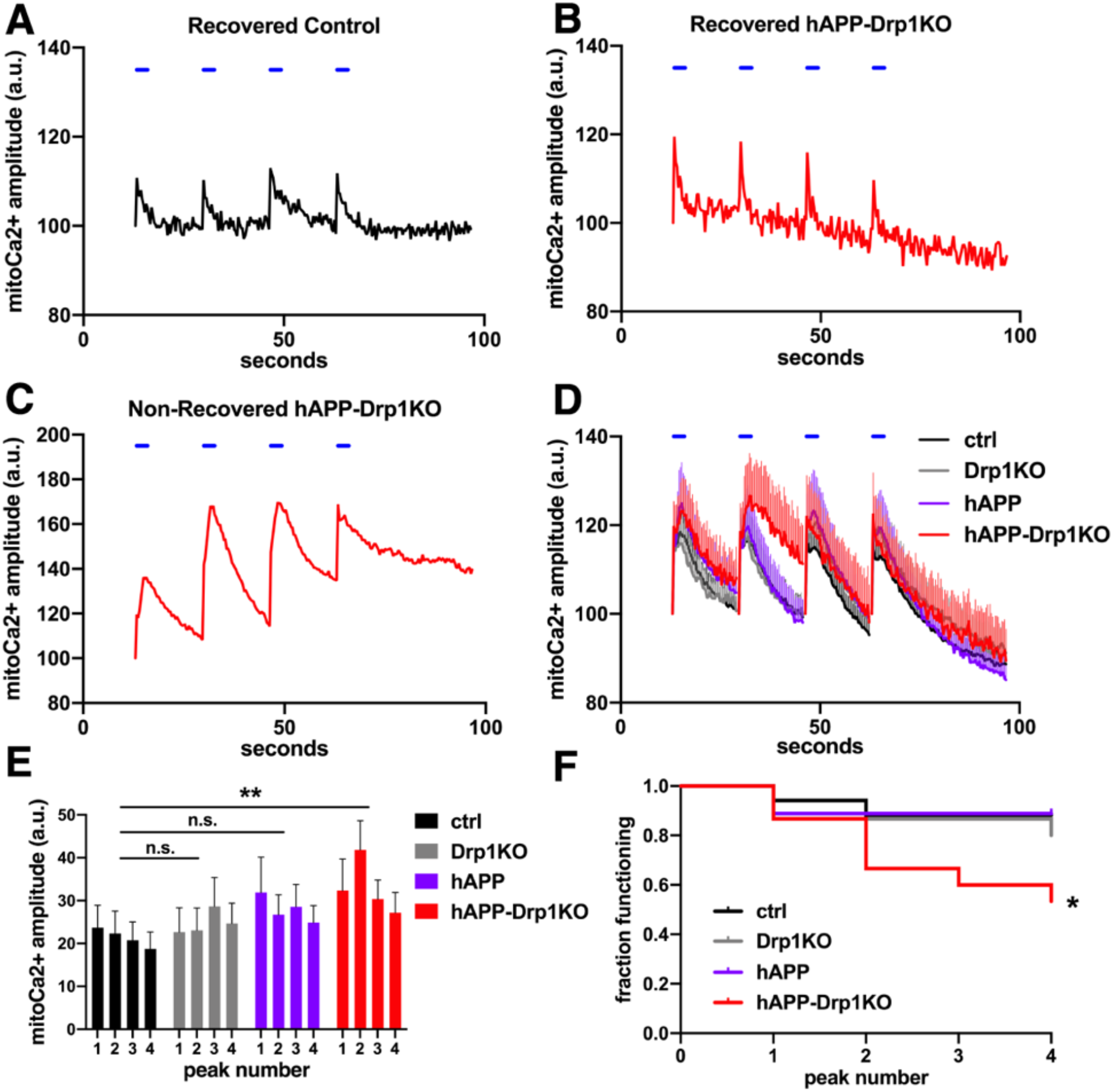
hAPP and Drp1KO combine to overload mitochondria with calcium. Primary hippocampal Drp1KO and control neurons were co-transfected with the mitochondrial calcium (mitoCa^2+^) sensor CEPIA3mt, mApple, and mutant hAPP or its control vector, and were subjected to a sequence of four individual electrical stimuli (30 Hz for 3s, blue horizontal lines) to evoke calcium entry. *A*, Example trace of a control neuron that recovers baseline mitoCa^2+^ levels following each stimulus. (*B*,*C*) Example traces of hAPP-Drp1KO neurons successfully (*B*) and unsuccessfully (*C*) recovering mitoCa^2+^ levels after evoked influx. *D*, Average mitoCa^2+^ levels for control (black), Drp1KO (grey), hAPP (purple) and hAPP-Drp1KO (red) neurons. *E*, Average amplitude for each mitoCa^2+^ peak in (*D*). The combination of hAPP and Drp 1KO, but neither of the perturbations alone, increased mitoCa^2+^ loading during electrically-evoked calcium entry compared to control. n= 15-18 coverslips/group (1 cell/coverslip), compilation of 6 independent experiments. Data show mean ± SEM; **p<0.01 by two-way repeated measures ANOVA and Holm-Sidak *post hoc* test. *F*, Graph shows the fraction of neurons from (*D*) that successfully recovered baseline mitoCa^2+^ levels following each of the four stimuli. The combination of hAPP and Drp1KO decreases the fraction of functional neurons; *p<0.05 by log-rank test.

### Drp1KO disrupts MAMs, but this does not underlie changes in mitoCa^2+^

We next investigated how hAPP and Drp1KO converge to disrupt mitoCa^2+^. We first focused on MAMs, which play a critical role in regulating calcium dynamics and transfer from the ER to the mitochondria (43). Interestingly, Aβ, mutant presenilins, and apoE4 can all increase the number, function and/or content of MAMs (44–46). In addition, Presenilin-1 and −2 (catalytic subunits of γ-secretase) and APP are all enriched in the MAM fraction of cells (47). These studies suggest a role for MAM dysfunction in AD pathogenesis. In addition, Drp1 function may be implicated in MAMs; although MAMs can form without Drp1 present (29), Drp1 is recruited to MAMs prior to mitochondrial fission (29).

Considering that Aβ has also been proposed to mediate toxicity by increasing Drp1 function (10), and given Drp1’s critical role in regulating mitochondrial morphology (19,22,42), we hypothesized that Drp1KO may prevent the ability of mutant APP to alter the number, structure or function of MAMs. We created three-dimensional reconstructions of confocal Z-stacks showing neuronal cell bodies from control and Drp1KO cells expressing EYFP targeted to the ER (EYFP-ER, yellow) and mito-FarRed (red), and identified contact regions as areas with persistent co-localization of the two probes for >3 min. Drp1KO neurons showed significantly fewer persistent MAMs, and smaller MAM area per mitochondrial content (Fig. 4, *A-E*), which was confirmed by electron microscopy (Fig. S4). However, hAPP did not alter the Drp1KO-induced decrease in the number and size of MAMs (Fig. 4, *A-E*). Therefore, the effects of Drp1KO on MAM formation are unlikely to underlie the synthetic effect of Drp1KO and hAPP on mitoCa^2+^.

**Figure 4.**
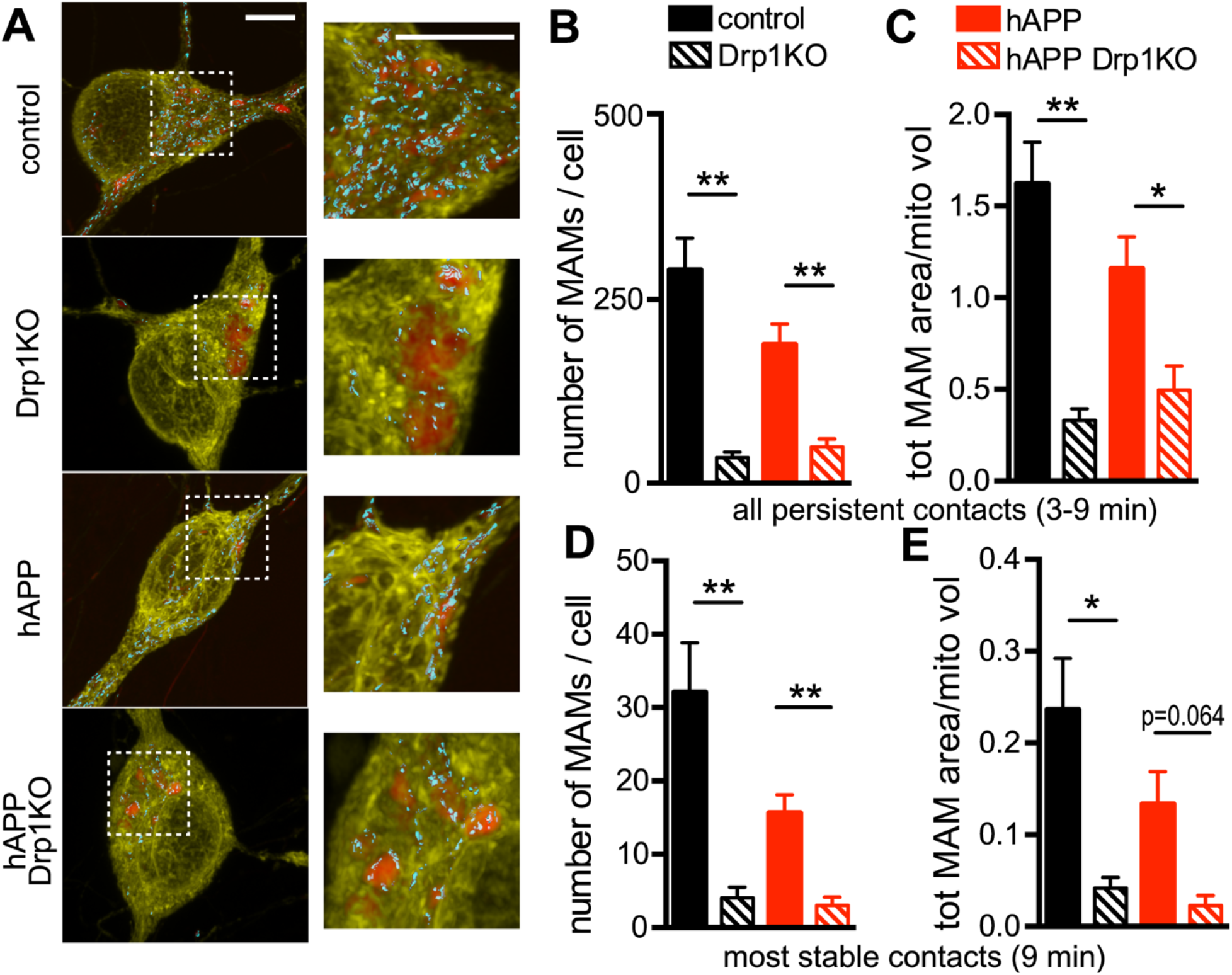
Drp1 loss decreases MAMs in cultured neurons. Hippocampal neurons from Drp1^lox/lox^ mice were co-transfected with Cre (to delete Drp1; Drp1KO), mutant hAPP, and/or control vector (control), as well as reporters to visualize the ER (yellow, eYFP-ER) and mitochondria (red, mitoFarRed). *A*, Threedimensional reconstructions of confocal Z-stacks (rendered via max projection) showing neuronal cell bodies with MAMs identified by areas showing ER-mitochondria co-localization (cyan; with surface rendering). *B*, In the presence or absence of hAPP, Drp1KO cells showed fewer persistent MAMs (defined as contacts lasting 3–5, 6–8, or 9 min) than Drp1WT cells. *C*, Drp1KO decreased the total area of ER-mitochondria contacts (normalized to total mitochondrial volume) with or without hAPP. (*D* and *E*) In the most stable contacts (lasting 9 min), both Drp1KO expression reduced MAM and total MAM area. *(B-E),* hAPP alone had no significant effect on MAMs as compared to control. n=8-11 coverslips/group (with 11–12 cells/group), compilation of 3 experiments. Data show mean ± SEM; p=0.064, *p<0.05, **p<0.01 by Welch’s ANOVA and Games-Howell *post hoc* test. Scale bars are 5 μm.

### Mitochondrial Ca^2+^ overload is not caused by excessive cytCa^2+^

Considering that Drp1KO disrupts MAMs, we asked whether the mitoCa^2+^ overload in hAPP-Drp1KO neurons is driven by excessive mitoCa^2+^ import from the cytosol, rather than the ER. To distinguish between an intrinsic increase in Ca^2+^ transport into mitochondria versus increased mitoCa^2+^ influx secondary to elevated cytCa^2+^, we examined cytCa^2+^ levels with GCaMP6f, a fluorescence-based calcium sensor with high temporal resolution (48). hAPP alone increased cytCa^2+^ versus control in response to electrical field stimulation (30hz for 3s and 10hz for 60s), consistent with prior work showing that mutant APP increases cytCa^2+^ levels following Ca^2+^ influx through the plasma membrane or from the ER (49). Drp1KO alone did not affect the extent of increase in cytCa^2+^ in response to electrical stimulation, however, the combination of hAPP and Drp1KO markedly decreased cytCa^2+^ (Fig. 5*A* and *B*). Although the mechanism and significance of this synergistic decrease in cytCa^2+^ is unclear, the low cytCa^2+^ in hAPP-Drp1KO neurons suggests that the increased mitoCa^2+^ in that genotype may result from increased shuttling of cytCa^2+^ into the mitochondria.

**Figure 5.**
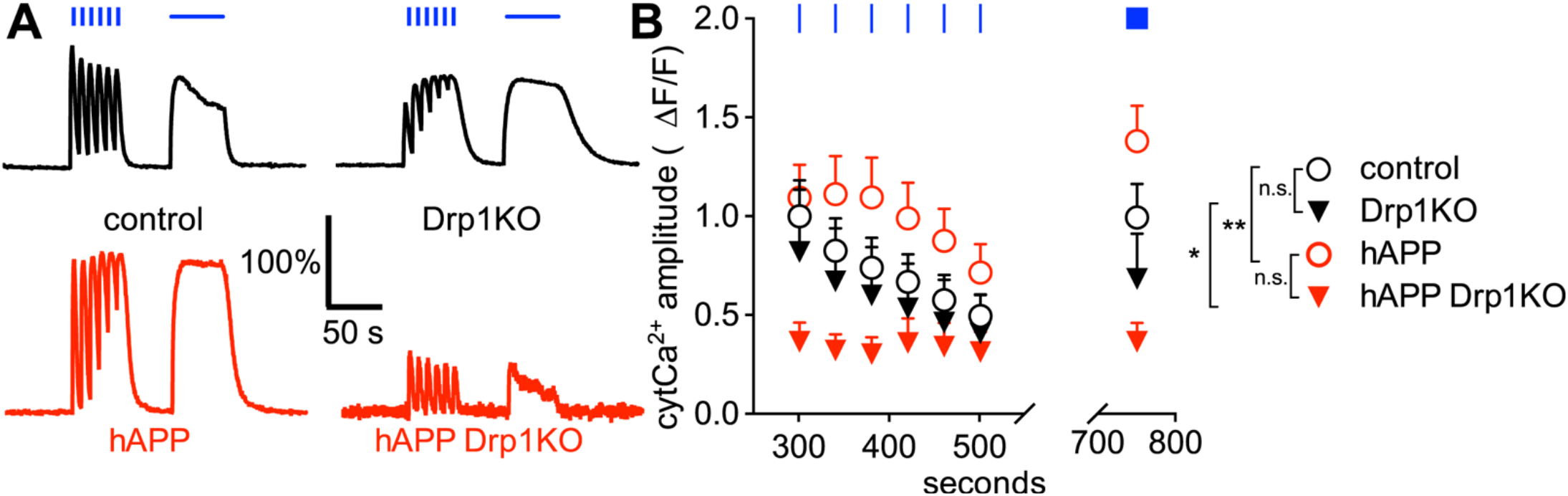
hAPP expression increases evoked cytosolic calcium in the cell body of neurons, but drastically decreases evoked calcium in the absence of Drp1. (*A* and *B*) Drp1KO and control neurons were co-transfected with the cytosolic calcium (cytCa^2+^) sensor GCaMP6f ***(81)*** and either mutant hAPP or vector control, and subjected to electrical stimulation (30 Hz for 3 s (blue vertical bars) and 10 Hz for 60 s (horizontal blue bar)). Drp1KO alone had no significant effect on the amplitude of evoked cytCa^2+^, whereas hAPP expression increased cytCa^2+^ in the presence of Drp1 and decreased cytCa^2+^ in the absence of Drp1. n=7-8 coverslips/group (with 17-60 cells/group), compilation of 3 experiments. Data are representative traces normalized to baseline and control (*A*) and means ± S.E.M. (*B*) *p<0.05 Drp1KO versus hAPP Drp1KO, **p<0.01 control versus hAPP by two-way ANOVA and Holm-Sidak test. Other comparisons (control vs. Drp1KO, hAPP vs. hAPP Drp1KO) were not significant (n.s.).

### mitoCa^2+^ overload occurs independent of an effect of hAPP on ATP levels

We previously showed that neurons lacking Drp1 have decreased mitochondrial-derived ATP levels (20). hAPP can also impair respiratory function (50,51), and optimal cytCa^2+^ and mitoCa^2+^ are required for efficient oxidative phosphorylation (37). To determine if hAPP converges with Drp1KO to produce energy failure, we co-transfected Drp1^lox/lox^ hippocampal neurons with a FRET-based ATP sensor (ATP1.03^YEMK^, (52)), and either Cre (to delete Drp1), hAPP, or vector control. To specifically measure mitochondrial-derived ATP we examined FRET levels in the acute absence of glucose and in the presence of glycolytic inhibitors 2-deoxyglucose (2DG) and iodoacetate (20,42). As expected, under these conditions Drp1KO axons could not maintain ATP levels (20), and ATP levels at the cell body were also slightly but significantly decreased (Fig. 6, *A* and *B* and Fig. S5, *A* and *B*). However, hAPP expression alone did not cause mitochondrial ATP deficits at the cell body or synapse, and hAPP did not alter ATP levels in Drp1KO neurons, even upon electrical stimulation to increase energy demands.

**Figure 6.**
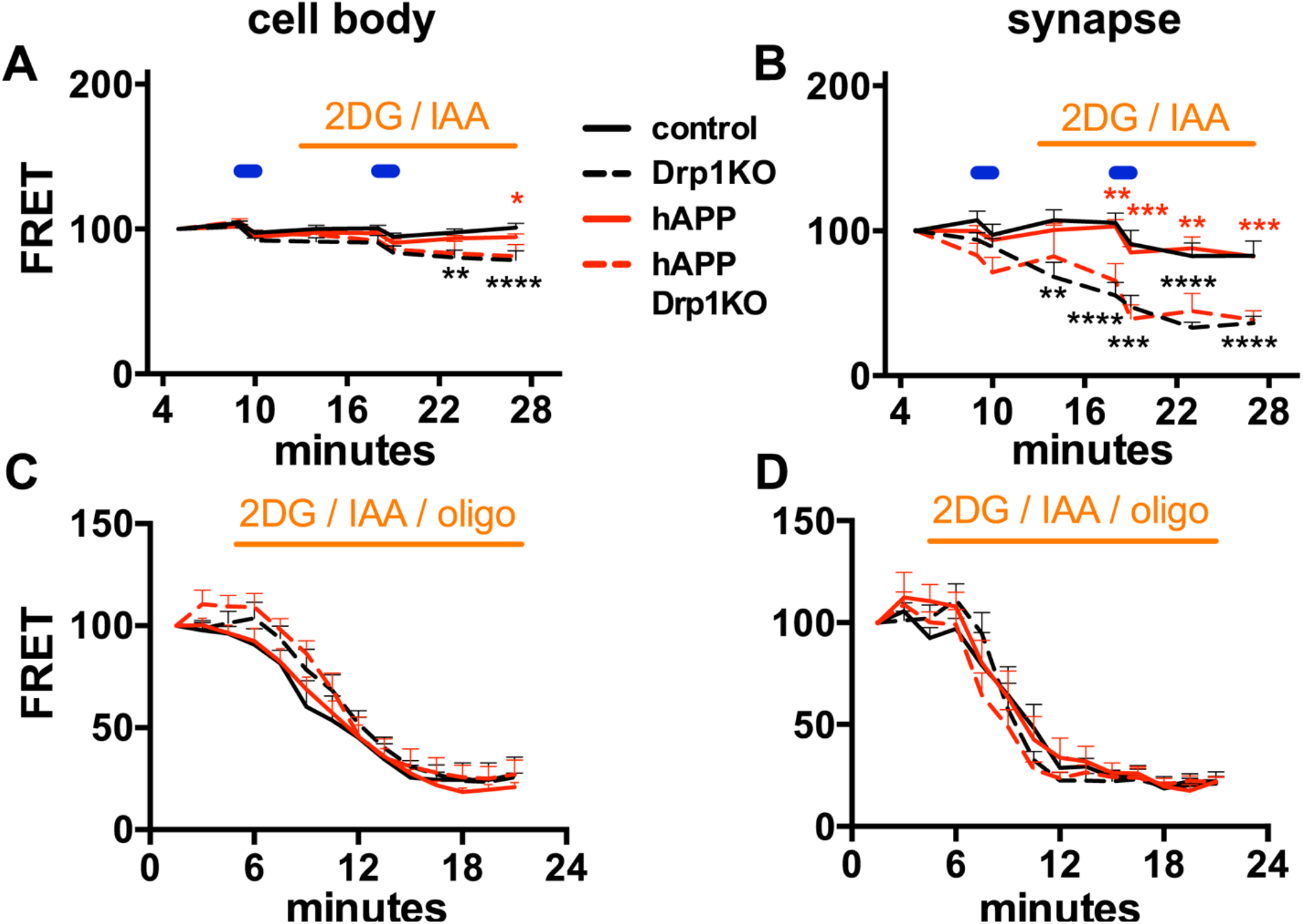
Drp1KO, but not hAPP expression, reduces mitochondria-derived ATP at synapses more than at cell bodies. Drp1KO and control neurons were co-transfected with an ATP-based FRET sensor (ATP1.03^YEMK^) ***(49)*** and either mutant hAPP or vector control. *A,* When forced to rely on mitochondria for ATP (acute absence of glucose, addition of glycolytic inhibitors 2-deoxyglucose (2DG) and iodoacetate (IAA); orange horizontal bar), Drp1KO neurons with or without hAPP had only slightly decreased ATP levels at the cell body after stimulation (10 Hz * 60 s, blue horizontal bars). *B,* In contrast, Drp1KO neurons with or without hAPP had markedly reduced ATP levels at the synapse under these conditions. hAPP did not affect ATP levels. n=6–12 coverslips/group (with 67–105 boutons and 15–22 cells per group), compilation of 4 experiments. *(C* and *D)* To estimate basal ATP consumption, we simultaneously blocked glycolytic production with 2DG and IAA and respiration with oligomycin (oligo). Rates of consumption did not differ across groups at the cell body (*C*) or the synapse (*D*), as indicated by the initial slope of decline in ATP level. n=6-8 coverslips/group (with 56–68 boutons and 6–10 cells per group), compilation of 3 experiments. Data are means ± S.E.M.; *p<0.05, **p<0.01, ***p<0.001, ****p<0.0001 control versus Drp1KO (black) and hAPP versus hAPP Drp1KO (red) by two-way ANOVA with repeated measures and Holm-Sidak test.

Given the energetic requirement needed to maintain calcium gradients, disruptions in calcium dynamics could also impact ATP consumption. To test this, we monitored the rate of decline in ATP levels after blocking all energy production using oligomycin (to inhibit respiration), 2DG and iodoacetate (to block glycolysis). However, neither Drp1KO nor hAPP affected the rate of ATP consumption (Fig. 6, *C* and *D*). Therefore, hAPP does not synergize with loss of Drp1 to disrupt mitochondrial-derived ATP in vitro. Instead, our data show that hAPP combines with a mitochondrial insult (here a change in mitochondrial fission) to disrupt mitochondrial Ca^2+^ homeostasis independent of energy levels, although this does not exclude the possibility that energy levels are ultimately affected in vivo.

## DISCUSSION

Changes in mitochondrial fission have been implicated in the pathophysiology of AD, but we understand little about how the level of fission influences the toxicity of key AD proteins. Here, we show that loss of fission alone does not alter either cytosolic or mitochondrial Ca^2+^ homeostasis, but it converges with mutant hAPP to produce mitochondrial Ca^2+^ overload, seemingly independent of Drp1KO’s effect of decreasing ATP levels in neurons.

### Drp1 protects mitochondria against hAPP-induced mitochondrial Ca^2+^ overload

AD and other neurodegenerative diseases are multifactorial, but little is known regarding how contributing processes converge to produce toxicity. Here, we dissect how two stressors implicated in the pathophysiology of AD combine to produce toxicity. First, we show that the dramatic, synergistic behavioral deficits produced by combining Drp1KO and hAPP *in vivo* cannot be attributed to changes in the mitochondrial morphology or content. This is surprising, since Drp1 has a primary role in regulating mitochondrial morphology (Berthet A et al. (2014), Shields et al. (2015)), and has been hypothesized to impact mitochondrial turnover (53). Instead, we show that loss of mitochondrial fission converges with hAPP to cause mitoCa^2+^ overload. Future studies are required to determine if hAPP predisposes to mitoCa^2+^ overload by increasing Aβ secretion from cells, or if other APP fragments may also contribute.

We hypothesized that the changes in mitochondrial Ca^2+^ might be explained by the effects of Drp1KO and hAPP on ER-mitochondria contacts. Indeed, Aβ, APOE4 and mutant presenilins can increase MAM formation (44–46), and Drp1 is recruited to MAMs (29). However, it has remained unclear if mitochondrial fission at these sites also plays a role in forming or maintaining these contacts. Here, we show that neurons require Drp1 to maintain MAMs, consistent with prior findings that inhibiting Drp1 decreases MAMs, while increasing mitochondrial fragmentation increases MAMs (54). The mechanism by which Drp1KO disrupts MAMs remains to be defined, but we hypothesize that steric factors contribute, especially given the characteristic swollen, rounded mitochondria at the cell body of Drp1KO neurons (19,20). However, hAPP failed to amplify the effect of Drp1KO on MAMs, suggesting that alteration in the number of MAMs is unlikely to underlie the hAPP-Drp1KO synergy. Instead, our data support that Drp1KO promotes mitoCa^2+^ overload in hAPP-Drp1KO neurons by driving excessive influx of cytosolic Ca^2+^ into mitochondria. We speculate that increased mitoCa^2+^ is driven by the changes in mitochondrial shape, which may disrupt the relative capacity for mitoCa^2+^ import versus export, as proposed in a recent study showing that Drp1KO myofibers also have increased mitoCa^2+^ influx following electrical stimulation (55). However, increased mitoCa^2+^ may also cause changes in mitochondrial shape and degeneration. Indeed, toxic insults that increase cytCa^2+^ typically produce more fragmented mitochondria indicative of increased mitochondrial fission (56), and increasing mitoCa^2+^ by overexpressing the mitochondrial Ca^2+^ uniporter (MCU) drives mitochondrial fragmentation in neurons (57), while inhibiting MCU normalizes mitoCa^2+^ and protects against degeneration(55). Although it remains unclear how exactly mitochondrial fragmentation influences toxicity, our findings raise the possibility that mitochondrial fission is a protective mechanism that enables mitochondria to either prevent or cope with the excessive mitoCa^2+^ influx.

### hAPP-Drp1KO mitochondrial Ca^2+^ overload occurs independent of ATP

Our dissection of the Aβ-Drp 1 interaction identified increases in mitoCa^2+^ that are seemingly independent of effects on mitochondrial-derived ATP. Although it is well recognized that mitochondria have functions such as Ca^2+^ homeostasis that are distinct from their roles in ATP production, these factors are often difficult to dissociate. Indeed, sufficient mitoCa^2+^ is required to maintain respiratory enzyme function (58,59), whereas excessive mitoCa^2+^ depolarizes mitochondria, transiently (60) or continuously, by opening the permeability transition pore (MPTP) (61). However, here we show that mitoCa^2+^ was disrupted without altering mitochondrial-derived ATP levels. Although ATP levels may ultimately change in vivo, this shows that changes in mitoCa^2+^ occur proximally, and may be the primary or initiating event. Sporadic AD is a multifactorial disorder, and this type of mechanistic insight is necessary to begin to understand how distinct toxic insults may combine to drive neuronal dysfunction and degeneration, and to provide insight into how such complicated interactions might be targeted therapeutically.

### Therapeutic potential of decreasing mitoCa^2+^ and Drp1 in AD

This work expands the field’s knowledge on the impact of Ca^2+^ homeostatic changes and mitochondrial function in models of AD. It is tempting to speculate that the same mitoCa^2+^ changes we observed in isolated neurons occur in vivo, and might contribute to the synergistic worsening of learning and memory in Drp1KO-hAPP mice. Indeed, mutations in APP and other proteins implicated in AD have been shown to disrupt cytosolic and ER calcium homeostasis (62–64), and Aß oligomers can increase mitoCa^2+^ levels, perhaps due to increased uptake secondary to increased expression of the mitoCa^2+^ uniporter (MCU) (65,66). Interestingly, patients with AD have decreased expression of the mitochondrial Na^+^/Ca^2+^ exchanger (NCLX). Moreover, 3xTg-AD mice have impaired mitoCa^2+^ efflux, and rescuing mitoCa^2+^ efflux by expressing NCLX markedly improved behavioral deficits and Aß and tau pathology (67). However, further work is required to fully understand the impact of mitochondrial function and calcium dysfunction in AD, and the possible therapeutic opportunities.

Lastly, our data suggest that Drp1 inhibition may be a risky target for AD therapeutics. While excessive Drp1 function has been hypothesized to mediate Aβ toxicity (32), Drp1KO not only fails to prevent hAPP toxicity, it actually markedly exacerbates it. While our findings are consistent with the possibility that partial Drp1 inhibition could be protective (2,12), careful calibration of the fission-fusion balance may be required for therapeutic efficacy. Our data clearly support the hypothesis that fission-fusion balance is necessary to support cellular function, and that an excessive shift in either direction can cause neuronal dysfunction (13–16,68–70).

## EXPERIMENTAL PROCEDURES

### Animals

Floxed Drp1 (22) and hAPP-J20 (34) mice have been described. CamKCre mice (71) were obtained from Jackson Laboratory. Mice were group-housed in a colony maintained with a standard 12 h light/dark cycle and given food and water ad libitum. Experiments were performed on age-matched mice of either sex. No differences between sexes were noted in any of the experiments. Experiments were conducted according to the *Guide for the Care and Use of Laboratory Animals,* as adopted by the National Institutes of Health, and with approval of the University of California, San Francisco, Institutional Animal Care and Use Committee.

### Behavioral testing

Learning and memory was assessed with the Morris water maze (MWM) test (72). Briefly, 12 sessions of visible platform training were performed prior to hidden platform training as a control. Subsequently, mice underwent two sessions of hidden-platform training separated by a 2 h intersession rest. Each session consisted of two trials. This training was performed each day for 7 days. The platform was removed and memory probe trials were performed 24 h and 72 h after the last training day.

EthoVision video-tracking system (Noldus, Netherlands) was used to record and track mice. Open field locomotor activity was performed as described (19). Briefly, mice were habituated for at least 1 h before recording activity for 15 min with an automated Flex-Field/Open Field Photobeam Activity System (San Diego Instruments, San Diego, CA). All behavioral experiments were performed with the examiner blinded to genotype.

### Histology and immunocytochemistry

For histology experiments, mice were anaesthetized and perfused with phosphate buffered saline (PBS) and then 4% paraformaldehyde (PFA). Brains were then removed, postfixed in PFA overnight, and cryoprotected in 30% sucrose. Coronal brain slices (30 μm) were prepared using a sliding microtome (Leica SM2000R).

For immunocytochemistry experiments, neuronal cultures were prepared as described below on coverslips and fixed in 4% PFA for 20 minutes.

For immunofluorescence, sections and coverslips were blocked for ≥1 h in PBS with 0.2% or 0.5% (for 82E1) Triton X-100 and 5-10% bovine calf serum and then incubated with primary antibodies overnight at RT. The following primary antibodies were used: chicken anti-MAP2 (1:1000; Abcam Cat# ab5392, RRID:AB_2138153); mouse anti-NeuN (1:1000; Millipore Cat# MAB377, RRID:AB_2298772); rabbit anti-Tom20 (1:500; Santa Cruz, Cat# SC-11415, RRID:AB_2207533); mouse anti-MAP2 (1:1000; Millipore Cat# MAB3418, RRID:AB_94856); rabbit anti-calbindin (1:20000; Swant Cat# 300, RRID:AB_10000347); mouse anti-8E5 (for human APP; 1:5000 (35)). Sections and coverslips were rinsed and incubated for 2 h at RT with the corresponding secondary antibodies: Alexa Fluor or DyLight 350, 488, 594, or 647 anti-mouse, chicken, or rabbit IgG (1:100–days before live imaging or a1:500; Invitrogen). For peroxidase staining, sections were quenched with 3% H_2_O_2_ and 10% methanol in PBS, and blocked in 10% bovine calf serum and 0.2% gelatin in PBS with 0.5% Triton X-100. They were incubated with mouse anti-82E1 (1:1000; IBL – America (Immuno-Biological Laboratories) Cat# 10326 RRID:AB_10705565), followed by biotinylated goat antimouse IgG (1:300; Vector Laboratories, Burlingame, CA; BA-1000, RRID:AB_2313606), and subsequently streptavidin-conjugated horseradish peroxidase (HRP) (1:300; Vectastain ABC kit, Vector Laboratories). Immunostaining was visualized with hydrogen peroxide and 3,3’-diaminobenzidine (DAB, Sigma).

Brain sections and coverslips were imaged with a laser-scanning confocal microscope (Zeiss LSM510-Meta, Zeiss LSM780-NLO FLIM, or Leica TCS SP8X) with a 63x (1.4 NA) PlanApo oil objective or (1.2 NA) C-Apochromat water objective, a Nikon Ti-E inverted microscope with a 60x (1.2 NA) PlanApo water objective, or a Keyence inverted microscope BZ-9000 with a 10x (0.45 NA) CFI PlanApo λ objective. Volume was calculated with the Cavalieri principle (73). Quantification of fluorescence and area were performed blind to genotype with MetaMorph software (version 7.7.3.0; Universal Imaging, RRID:SciRes_000136). Neuronal density was calculated by dividing the total fluorescence of NeuN in 100μm^2^ by the average NeuN intensity per CA1 neuron. Quantification of cells with swollen mitochondria was scored blind to genotype, based on the presence of 3 or more swollen mitochondria in a cell (a subjective criterion chosen to distinguish Drp1cKO versus control mitochondria). Amyloid plaque load was calculated based on % area of the hippocampus covered by plaques. Colocalization of MAM images was analyzed using Imaris software and the Surface-Surface colocalization extension.

### Neuronal culture and live imaging

Postnatal hippocampal neuronal cultures were prepared from P0 Drp1^lox/lox^ mice as described (19) and transfected via electroporation (Amaxa) with one or more of the following constructs, all expressed in the pCAGGS vector downstream of the chicken actin promoter (74): ATP-YEMK (kind gift of Dr. Noji, Osaka University) (52), mCherry-synaptophysin (75), Cre recombinase (19), hAPP mutant (Swedish, Indiana) (39,40), ires-mApple, mitoGFP (5), GCaMP6f (48), Cepia3mt(41), ER-eYFP (Clontech), or mitoFarRed. mitoFarRed was generated by fusing TagRFP657 (kind gift from Vladislav Verkhusha (Albert Einstein)) to the mitochondria-targeting sequence, cytochrome C oxidase subunit VIII (76,77). Neurons were cultured for 8–11 days before live imaging or analysis.

Live imaging was performed in Tyrode’s medium (pH 7.4; 127 mM NaCl, 10 mM HEPES-NaOH, 2.5 mM KCl, 2 mM MgCl_2_, 2 mM CaCl_2_, with or without 30 mM glucose and/or 10 mM pyruvate) on a Nikon Ti-E inverted microscope with an iXon EMCCD camera (Andor Technology) and a perfusion valve control system (VC-8, Warner Instruments) controlled by MetaMorph Software. Live imaging for MAM studies was performed on a Zeiss LSM880 confocal microscope with Airyscan detector. Field stimulations (10 Hz*60 s and 30 Hz*3 s) were performed with an A385 current isolator and a SYS-A310 accupulser signal generator (World Precision Instruments). Glycolysis was inhibited with 2-DG [5 mM, Sigma-Aldrich] and iodoacetate [1mM, Sigma-Aldrich]. Respiration was inhibited with oligomycin [3μM].

For GCaMP6f and Cepia3mt calcium experiments, images were obtained (490/20 ex, 535/50 em, Chroma) every 200 msec. A region of interest was drawn over the cell body, excluding the nucleus, and the background-subtracted fluorescence was calculated for each timepoint, normalized to the baseline level of background-subtracted fluorescence and control.

For FRET experiments, sequential images were taken in the CFP (430/24 ex, 470/24 em), YFP (500/20 ex, 535/30 em), and FRET channels (430/24 ex, 535/30 em) with an ET ECFP/EYFP filter set (Chroma). Synaptic boutons were identified based on morphology. The FRET/donor ratio was calculated for each bouton and cell body as described (78), where FRET = (I_FRETCFP_*BT_CFP_ – I_YFP_*BT_YFP_) / I_CFP_, such that I_X_ is the background-corrected fluorescence intensity measured in a given channel. BT_CFP_ (donor bleed through) and BT_YFP_ (direct excitation of the acceptor) were calculated by expressing CFP and YFP individually and determining the ratios of I_FRET_/I_CFP_ and I_FRET_/I_YFP_, respectively.

### Electron microscopy

For EM of hippocampal neurons, DIV10 cultures were fixed in 4% PFA for 2 hours, and then incubated with 3% glutaraldehyde and 1% PFA in 0.1 M sodium cacodylate buffer (pH 7.4) overnight. Following the fixation, the cultures were processed through 2% osmium tetroxide and 4% uranyl acetate, then dehydrated and embedded in Eponate 12 resin (Ted Pella Inc., Redding, CA). Ultra-thin sections were cut at 1-μm thick, collected on copper grids, and imaged in a Phillips Tecnai10 transmission electron microscope using FEI software. Quantitative analysis was performed on digital EM images obtained with a charge coupled device (CCD) camera at a final magnification of 11500 (Berthet et al., 2014). The quantification of EM samples from cultured mouse hippocampal neurons was performed with the examiner blinded to the genotypes using MetaMorph software (version 7.7.3.0; Universal Imaging, RRID: SciRes_000136).

### Statistical analysis of Morris water maze

Morris Water Maze data has several characteristics that make longitudinal analysis complex: the data typically contain censoring, as the mice are removed from the water after a fixed amount of time if they fail to complete the task, the learning effect is often very non-linear; as healthy mice often learn the maze as well as they can before the last trial and thus stop systematically improving, there is typically a learning effect of both days of trials and number of trials given that day, which leads to a “saw-tooth” learning effect, and finally, a mean-variance relation is expected.

Rather than attempt to build a very complex statistical model to account for these data features, a summary measure analysis (79) was created, which greatly reduced the dimensionality of the problem and allowed for simple, robust, powerful and easily interpreted results. To do this, at each trial, each mouse is ranked (i.e., which mouse finished first, second, etc.). Mice that failed to locate the platform are considered “tied for last”. For each mouse, the average rank across all trials is then calculated. This simple composite score is used in standard analyses.

The outcomes considered were the average ranks of latency per mouse during hidden and visible trials. The data were fit to two linear mixed effects models (80) corresponding to these outcomes and to the factor genotype using the R package lme4 (RRID:nif-0000-10474) (81). Random effects for the effect of cohort on genotype were included, and the overall effect of genotype on average rank latency hidden and average rank latency visible tested using the Wald Chi-square test.

The fitted model was used to obtained estimates of the mean difference in ranks. The function sim() from the arm package (82) yielded 50,000 draws of the group effects, from which 95% confidence interval (CI) around each estimate as the 2.5^th^ and 97.5^th^ quantiles were calculated. p-values for differences between groups were calculated using the simulated differences. The p-values corresponding to the composite scores were not adjusted as the gatekeeping testing approach was appropriate to use here (83,84). The p-values corresponding to the rest of the outcomes were corrected for multiple comparisons using the method of Holm (85).

For the memory probe trials, two different outcomes were analyzed: latency to first target platform crossing and number of platform crossings. Data were fit onto a Cox proportional hazards regression model of latency on genotype using the R package survival (86). The proportional hazards assumption was examined by visual inspection of the curves of the natural logarithm of the cumulative hazard function versus latency for each of the four genotypes. The four curves were approximately parallel, indicating that the proportional hazards assumption was met for this dataset. Kaplan Meier nonparametric test was also performed to investigate the survival function for each genotype. Random effects for the effect of cohort on genotype were included.

The number of platform crossings by genotype data were fit to a Quasi-Poisson generalized linear model. The Quasi-Poisson generalized linear model accounts for overdispersion, allowing for more robust estimation. Two Quasi-Poisson models were used, one with an interaction term between genotype and cohort, to ensure that the treatment effect was relatively constant across cohorts, and the primary model was fit with only genotype alone, (assuming the effect of genotype was consistent across cohorts). A Deviance test was used to compare these two models, revealing no significant difference between them (p-value=0.166), implying that the effects were consistent across cohorts. Therefore, the model with only the effect of genotype was used.

Estimates of the relative risk of reaching the platform (RR), the 95% confidence interval of the relative risk, and the corresponding p-values were obtained from the Cox proportional hazards regression model. The mean difference in number of platform crossing, the 95% confidence interval of the difference, and the corresponding p-values were obtained from the Quasi-Poisson model. The p-values corresponding to these outcomes were corrected for multiple comparisons using the method of Holm (85).

## DATA AVAILABILITY

Data described in the manuscript is available upon request. Please contact Ken Nakamura: Ken Nakamura, MD, PhD, Gladstone Institute of Neurological Disease, 1650 Owens Street, San Francisco, CA 94158, Phone: (415) 734-2550; Fax: (415) 355-0824;

## ACKNOWLEDGMENTS

We thank Lennart Mucke for providing hAPP-J20 mice and for comments on the manuscript, Gladstone’s Behavioral Core and Histology and Light Microscopy Core for technical assistance, Ivy Hsieh and Eric Huang for help with electron microscopy, Jonathan Levy for guidance preparing lentivirus, Grisell Diaz-Ramirez, Clifford Anderson-Bergmant and Reuben Thomas for assistance with statistics, Jeff Simms for feedback on the manuscript, and Kathryn Claiborn for helping edit the manuscript.

## AUTHOR CONTRIBUTIONS

LS, HL, TMG and KNakamura designed research; LS, HL, KNguyen, HK performed research; ZD, TMG, DH, KAV, and MC provided guidance or reagents for critical techniques; LS, HL, KNguyen, ZD, TMG and KNakamura analyzed data; LS and KNakamura wrote the paper with assistance from co-authors.

## FUNDING AND ADDITIONAL INFORMATION

KN, LS, HL, KNguyen, HK, DH and ZD were supported by NIH RO1NS091902 and RF1AG064170 to KN. This work was also supported by NIH 5P30 NS069496 and NIH RR18928 to Gladstone Institutes.

## ABBREVIATIONS AND NOMENCLATURE

(2DG): 2-deoxyglucose
(PFA): 4% paraformaldehyde
(AD): Alzheimer’s disease
(Aβ): amyloid beta
(cytCa^2+^): cytosolic calcium
(CCD): charge coupled device
(Drp1): dynamin-related protein 1
(EYFP-ER): EYFP targeted to the ER
(Drp1cKO): hAPP mice that lack Drp1 in the CA1 and other forebrain neurons
(hAPP): human amyloid precursor protein
(MAMs): mitochondria-associated ER membranes
(mitoCa^2+^): mitochondrial Ca^2+^
(NCLX): mitochondrial Na^+^/Ca^2+^ exchanger
(MWM): Morris water maze
(MPTP): mitochondrial permeability transition pore

